# Genomics Characterization of an engineered *Corynebacterium glutamicum* in Bioreactor Cultivation under Ionic Liquid Stress

**DOI:** 10.1101/2021.09.29.462453

**Authors:** Deepanwita Banerjee, Thomas Eng, Yusuke Sasaki, Aparajitha Srinivasan, Asun Oka, Robin A. Herbert, Jessica Trinh, Vasanth R. Singan, Ning Sun, Dan Putnam, Corinne D. Scown, Blake Simmons, Aindrila Mukhopadhyay

## Abstract

*Corynebacterium glutamicum* is an ideal microbial chassis for the production of valuable bioproducts including amino acids and next-generation biofuels. Here we resequence engineered isopentenol (IP) producing *C. glutamicum* BRC-JBEI 1.1.2 strain and assess differential transcriptional profiles using RNA sequencing under industrially relevant conditions including scale transition and compare the presence *vs*. absence of an ionic liquid, cholinium lysinate ([Ch][Lys]). Analysis of the scale transition from shake flask to bioreactor with transcriptomics identified a distinct pattern of metabolic and regulatory responses needed for growth in this industrial format. These differential changes in gene expression corroborate altered accumulation of organic acids and bioproducts, including succinate, acetate, and acetoin that occur when cells are grown in the presence of 50mM [Ch][Lys] in the stirred-tank reactor. This new genome assembly and differential expression analysis of cells grown in a stirred tank bioreactor clarify the cell response of a *C. glutamicum* strain engineered to produce IP.

## 1. Introduction

Due to process advantages, biological methods for the production of amino acids over chemical synthesis methods fostered the identification of natural glutamine overproducing microbes (Kinoshita et al., 1958). Since then, *Corynebacterium glutamicum* has been used successfully to produce specialty glutamine and specialty amino acids to meet global demand. The advent of accessible whole-genome sequencing and mutagenesis methods have enabled researchers a clearer understanding of how specific isolates can overproduce these desired molecules, as well as how they have maintained productivity across geometrically-larger scales (Wolf et al., 2021; Pérez-García and Wendisch, 2018; Becker et al., 2018). Using *C. glutamicum* to produce non-native metabolites as next-generation biofuels is an attractive large-volume market with the potential to reduce global carbon emissions. Potential biofuels can be produced from terpenes, which use different metabolic precursors (reviewed in (Pérez-García and Wendisch, 2018)). We have previously described the heterologous expression of the terpenoid isopentenol (IP; also known as 3-methyl-3-buten-1-ol or isoprenol) pathway in *C. glutamicum* (Sasaki et al., 2019). Isopentenol can be used directly as a drop-in biogasoline (Chou and Keasling, 2012; S-CoA, 2008) or as a precursor to a jet fuel, DMCO (Baral et al., 2021). Producing IP was improved by the use of optimal pathway homologs, specific media formulation and aeration conditions and an empirically determined carbon/nitrogen ratio.

In this study we build upon this established system to analyze the behavior of *C. glutamicum* strains engineered to produce IP in a bioreactor. The bioreactor cultivation and process conditions can provide key diagnostic information essential to build robust production platform strains (Wehrs et al., 2019). In addition, it is also valuable to have an understanding of microbial response to the carbon feedstock that is anticipated for actual production. Here, we explore the use of plant-based lignocellulosic hydrolysate generated using ionic liquid (IL) as a pretreatment reagent. Toxicity from residual pretreatment reagents such as ILs is a known source of growth impediment (Hou et al., 2013; Santos et al., 2014). *C. glutamicum* is tolerant to many ILs, another attribute that makes it an ideal host for biomass conversion (Sasaki et al., 2019). In this study, we characterize an IP producing engineered *C. glutamicum* strain with long-read PacBio whole-genome sequencing. This high-quality assembly allowed accurate mapping for differential RNA expression analysis from a diagnostic fed-batch *C. glutamicum* IP production run. These side-by-side experiments characterize the cellular response to the IL, cholinium lysinate ([Ch][Lys]), when grown in a fed-batch stirred tank bioreactor.

## 2. Results

### 2.1. Characterization of Isopentenol Production and Ionic Liquid Tolerance in C. glutamicum Strains

We established that the strain reported in Sasaki *et al* 2019, *C. glutamicum* (previously referred to as ATCC 13032 NHRI 1.1.2) outperformed another isolate, ATCC 13032 Δ*cglIM* Δ*cgLIR* Δ*cgLIIR* (referred to as *“Δmrr”*) **(Figure 1A)**. *C. glutamicum* Δ*mrr* was first described in Baumgart *et al* 2013 and is a methylation-deficient strain widely used due to its improved plasmid transformation and genomic integration rate (Schäfer et al., 1997; Baumgart et al., 2013). When *C. glutamicum* BRC-JBEI 1.1.2 is used in conjunction with an IP production pathway, it can produce 300 mg/L IP from pure glucose, but the product titers are near the lower detection limit by GC-FID in the *C. glutamicum* ATCC 13032 Δ*mrr* strain. While only *C. glutamicum* BRC-JBEI 1.1.2 produced IP, both the type strain and this specific isolate tolerate high concentrations of exogenous ILs **(Figure 1B)**, suggesting that IL tolerance was a shared feature due to the cell membrane structure between these two isolates even if the available metabolic flux towards IP was different.

**Figure 1.**
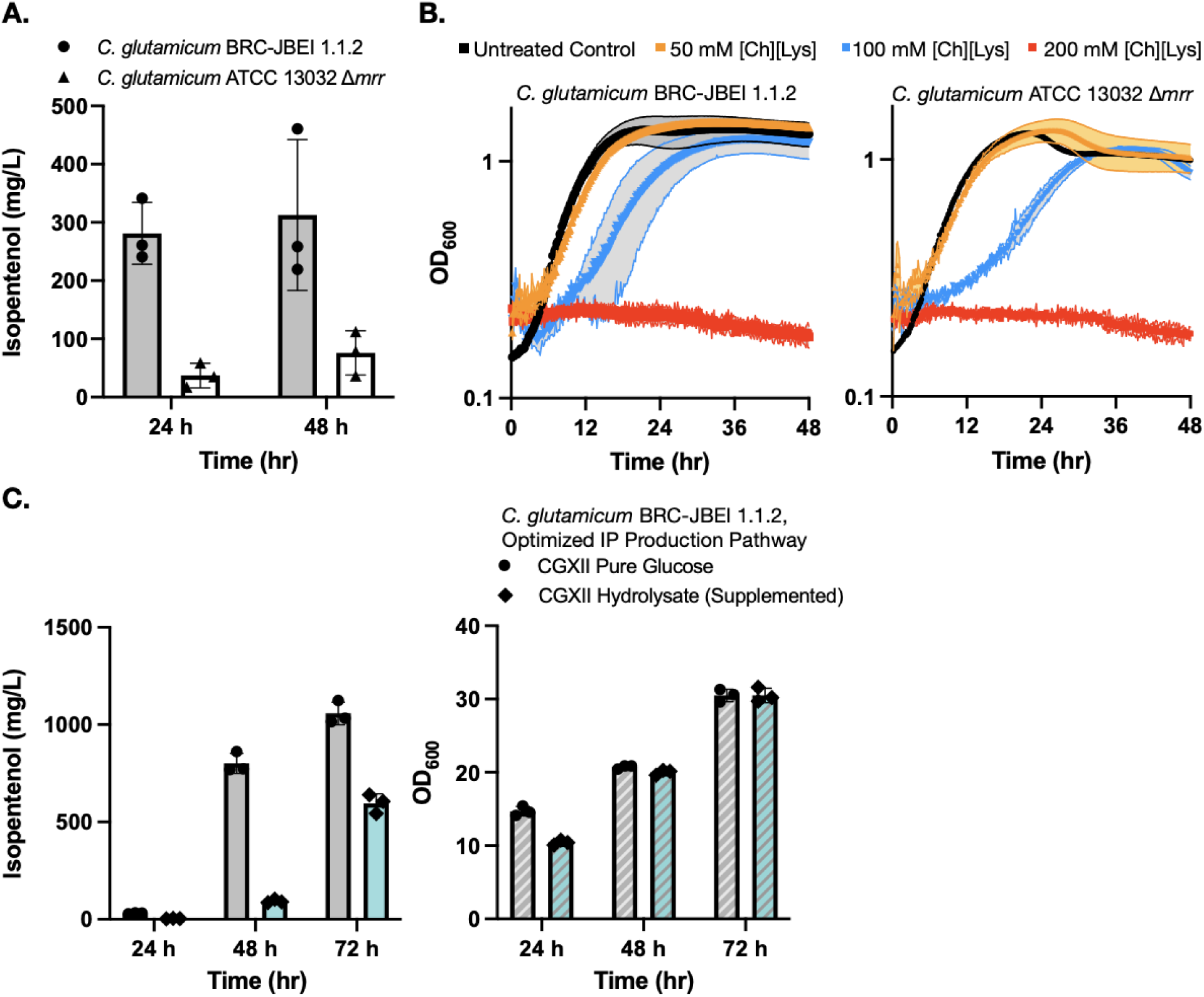
Growth and isopentenol production characterization of two genetically distinct engineered *C. glutamicum* strains. **(A)** Isopentenol (IP) production in *C. glutamicum* strains of the genotypes indicated harboring an IP production plasmid. Cells were cultivated in 24-well deep well plates. Isopentenol titers reported at 48-hour time points are corrected for evaporation in this plate format (Materials and Methods). **(B)** Growth curves for *C. glutamicum* strains of the indicated strain backgrounds cultivated in CGXII media in the presence or absence of the IL, cholinium lysinate ([Ch][Lys]). [Ch][Lys] was exogenously added to the culture media at the start of the time course. **(C)** Production of IP from *C. glutamium* grown in CGXII minimal media with pure glucose (4% w/v) or ensiled [Ch][Lys] pretreated sorghum hydrolysate. An optimized IP production plasmid carrying a *hmgR* variant from *Silicibacter pomeroyi* was used. The optical density of cultures as a proxy for cell density is noted on the right-hand panel.

We also confirmed the ability of *C. glutamicum* BRC-JBEI 1.1.2 to handle renewable carbon streams from sorghum biomass using an improved carbon extraction protocol enhanced by the use of ensiled biomass (Magurudeniya et al., 2021). The ensiling process enables naturally occurring lactic-acid secreting bacteria to partially decompose the hemicellulose in sorghum while stored in a silo before downstream processing. After ensiling, the biomass was pretreated with [Ch][Lys] followed by enzymatic saccharification (Materials and Methods). This hydrolysate contained 48.7 g/L glucose, 17.9 g/L xylose, and trace concentrations of aromatic compounds. Our optimized *C. glutamicum* BRC-JBEI 1.1.2 with an optimized IP production system had no detected growth defects when grown with 58% (v/v) hydrolysate supplemented media and produced 1 g/L IP from pure glucose or ∼600 mg/L IP from sorghum hydrolysate **(Figure 1C)**. These results showcase its versatility with handling real-world plant biomass derived carbon streams. For the remainder of this study, we focus on characterizing the genetic differences present in *C. glutamicum* BRC-JBEI 1.1.2 relative to other closely related *C. glutamicum* strains that might explain the IP production values between these two strains.

### 2.2 Genomic Characterization of C. glutamicum BRC-JBEI 1.1.2

16S rDNA sequencing (Hahne et al., 2018) confirmed the *C. glutamicum* Δ*mrr* strain as in the *C. glutamicum* ATCC 13032 strain background, but this same method indicated that *C. glutamicum* BRC-JBEI 1.1.2 was genetically closer to *C. glutamicum* CICC10112 or SCgG1/SCgG2. Only SCgG1 and SCgG2 have been characterized with whole-genome sequencing, and to our knowledge there was no additional information about *C. glutamicum* CICC10112 beyond the partial 16S ribosomal sequence. As 16S rDNA was not conclusive, we reasoned that the whole-genome sequencing in this IP producing strain would ensure an accurate reference genome in downstream RNAseq analysis if the improved performance observed in this strain was due to variants in the strain background. One of the major limitations in short-read sequencing is the difficulty in assembling overlapping contigs to generate a high-quality *de novo* assembly of a single contiguous read. Therefore, we chose Pac-Bio long-read sequencing (Koren and Phillippy, 2015) for optimal coverage over short read sequencing as a potential solution. However, routine methods for lysing and isolating *C. glutamicum* genomic DNA were insufficient for building high-quality genome assemblies since the physical lysis method we employed (Eng et al., 2018) shears DNA to fragments ranging from 2-8 kb in size. Detergent-based lysis methods failed to extract genomic DNA, even with prolonged incubation times. We developed a method to isolate larger DNA fragments approximately 20kb in size for the PacBio Sequel assembly pipeline using a Zymolyase protease treatment for cell lysis (see Materials and Methods). This modified DNA extraction protocol enabled us to use PacBio long read sequencing to generate a high-quality *de novo* genome assembly.

We now report a new genome assembly of a single contiguous scaffold of 3,352,276 bases with 53.83% GC content **(Figure 2)**. Genome-wide average nucleotide identity (ANI) confirmed this isolate was 99.9987% identical to *C. glutamicum* SCgG1 and SCgG2 as well as another sequenced *C. glutamicum* isolate, Z188. The average nucleotide identity alignment for the 28 sequenced *C. glutamicum* isolates has been deposited at the database of the Joint Genome Institute and is also included in **Supplementary Table S1**. *C. glutamicum* BRC-JBEI 1.1.2 differs from SCgG1 only by a few single nucleotide polymorphisms (∼10) and two additional genes that are absent from SCgG1, a putative transposase and a hypothetical protein coding sequence that is 414 bp in length. When *C. glutamicum* BRC-JBEI 1.1.2 was compared with more commonly used reference strains, *C. glutamicum* R and 13032 (Bielefeld), we identified genomic islands encoding genes unique to BRC-JBEI 1.1.2. Genome topology analysis also identified a 140 kb inversion in the genome of BRC-JBEI 1.1.2 isolate **(Figure 2A)**. Out of 3,097 genes, homology mapping indicated that 85% (2,641 genes) were at least 80% identical to known genes in *C. glutamicum* ATCC 13032. With a less restrictive % identity threshold of 50%, the identical ratio could account for 89% (2,777 genes). Nonetheless, 320 genes did not meet the minimum % identity threshold and could not be annotated with this reference genome **(Supplementary Figure S1)**.

**Figure 2:**
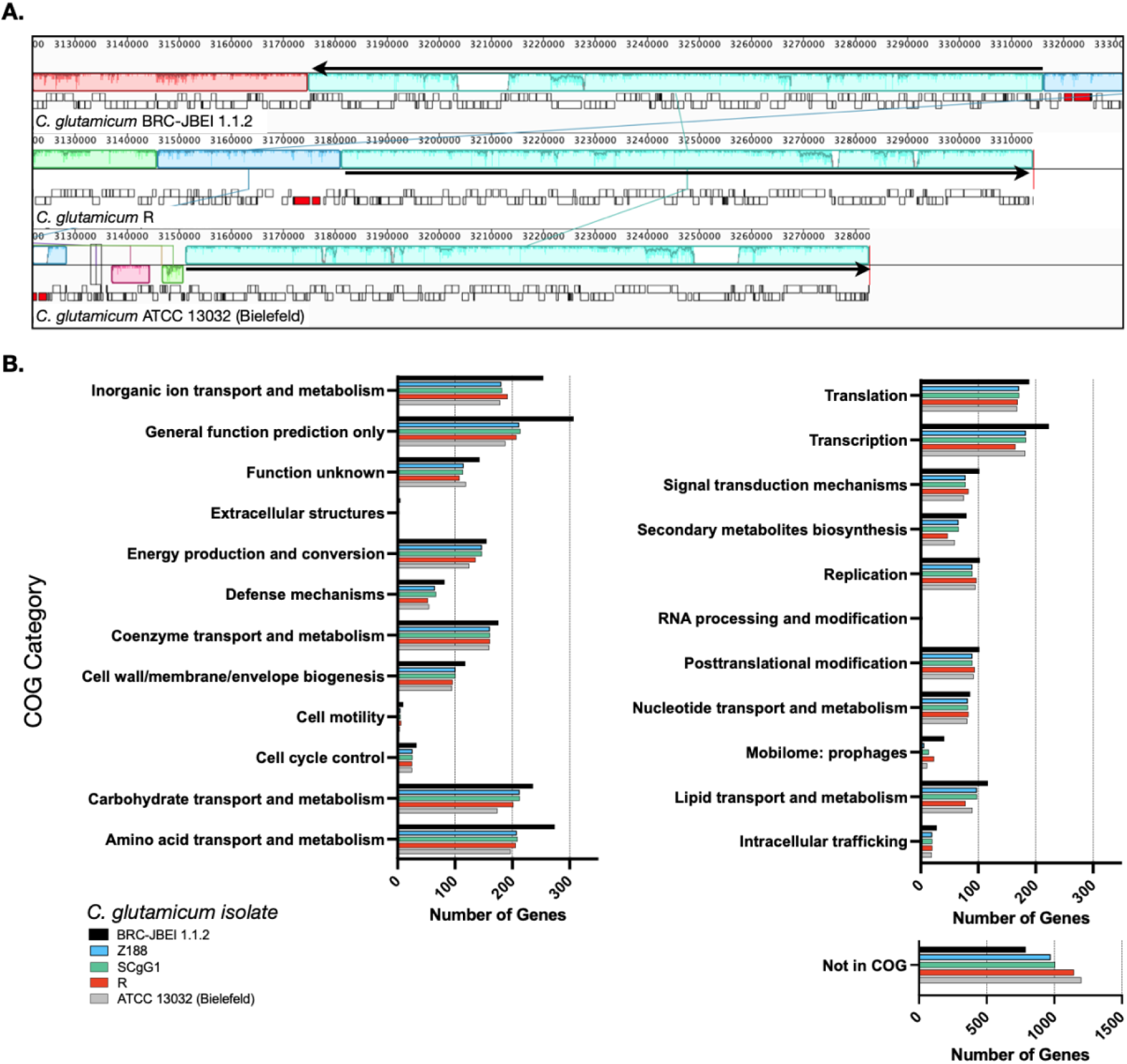
Comparison of the *C. glutamicum* BRC-JBEI 1.1.2 strain with closely related *C. glutamicum* strains. **(A)** A meta-analysis of gene function using clusters of orthologous genes (COGs) analysis. The total number of genes in each category for each strain is represented with colored bars as indicated. **(B)** Mauve genome alignment of *C. glutamicum* BRC-JBEI 1.1.2 with *C. glutamicum* R and 13032 (Bielefeld). Similar genomic regions share the same color across the 3 different genomes compared. A 140 kb chromosomal inversion is highlighted in light blue, and the relative direction of the inversion in each strain is indicated with a black arrow. Individual genes are indicated with open rectangles underneath the colored area.

Some of these unknown genes that were unique to BRC-JBEI 1.1.2 might be related to the catabolism of IL. Intriguingly, a putative choline dehydrogenase, *Ga0373873_2846*, showed only 40% identity to other known choline dehydrogenases primarily found in gram-negative microbes such as *Burkholderia phytofirmans* PsJN and *Cupriavidus basilensis* FW507-4G11. Meta-COG analysis of these four *C. glutamicum* genomes revealed that *C. glutamicum* BRC-JBEI 1.1.2 contains over 100 additional genes related to the transport or metabolism of inorganic ions, carbohydrates, and amino acids, suggesting a broader metabolic capacity to utilize a more significant number of substrates than the type strain **(Figure 2)**. In summary, this genome sequencing analysis was valuable for characterizing differences between *C. glutamicum* BRC-JBEI 1.1.2 and the more intensely studied type strain ATCC 13032. Due to its similarity with SCgG1 and SCgG2, *C. glutamicum* BRC-JBEI 1.1.2 is likely an industrial glutamate overproducing isolate but has more annotated transport and metabolic systems than its nearest neighbors, SCgG1, SCgG2, and Z188 that need further characterization.

### 2.3. Transcriptome Analysis Identifies Changes in C. glutamicum Metabolism on Scale-up

Next, we sought to build a systems-level understanding of *C. glutamicum* gene expression changes in bioreactors upon exogenous ionic liquid treatment. This data could be useful for subsequent Design-Build-Test-Learn (DBTL) cycles in providing the diagnostic information for future strain optimization strategies (Opgenorth et al., 2019). We prepared samples from sequential time points during a scaleup campaign to analyze shifts in gene expression as a proxy for changes in metabolic and regulatory behavior in both [Ch][Lys] treated and untreated runs. First we determined if the failure to produce IP was due to loss of the production pathway, possibly due to loss of the plasmid-borne IP pathway genes. The IP production pathway is composed of 5 genes in 2 adjacent operons under the *trc* and *lacUV5* promoters, namely *mk, pmd* and *atoB, hmgS, hmgR respectively*. Using the transcripts per million (TPM) metric, we examined absolute gene expression levels as well as changes over the course of the production campaign. The IP pathway started off high for both *hmgR* and *hmgS* in the shake flask (200,000 TPM), but expression of these two genes decreased between 10-16 x over the duration of the 65 hour fed-batch. Expression amounts of *atoB* in the shake flask were comparatively lower (1,500 TPM) but decreased 4 x at the shake flask to bioreactor transition. *atoB* TPM counts remained low for the duration of the subsequent time points. Since the pathway genes were still expressed during this run, we then focused on analyzing gene expression changes in the native *C. glutamicum* genome.

To interpret the differential gene expression results with genes identified in the new assembly for *C. glutamicum* BRC-JBEI 1.1.2, we mapped gene names and identifiers from *C. glutamicum* ATCC 13032 back onto the open reading frames (ORFs) in *C. glutamicum* BRC-JBEI 1.1.2 as genes in the type strain genome have been broadly characterized. We used a medium confidence cutoff of 70% identity to capture most homologs when analyzing this dataset. First, we characterised gene expression upon inoculating cells from the seed culture in a shake flask to the bioreactor. This differential gene expression (DEG) was calculated as the ratio of an early time point in the bioreactor (6.5 hours post inoculation in the stirred tank) divided by values from the seed culture immediately before transfer. This time point was chosen to give cells approximately three doublings to ensure the cells were rapidly growing under these new conditions. The result showed differential expression of 258 genes after 6.5 hours **(Figure 3**, and **Supplementary Data, Dataset S1)**.

**Figure 3.**
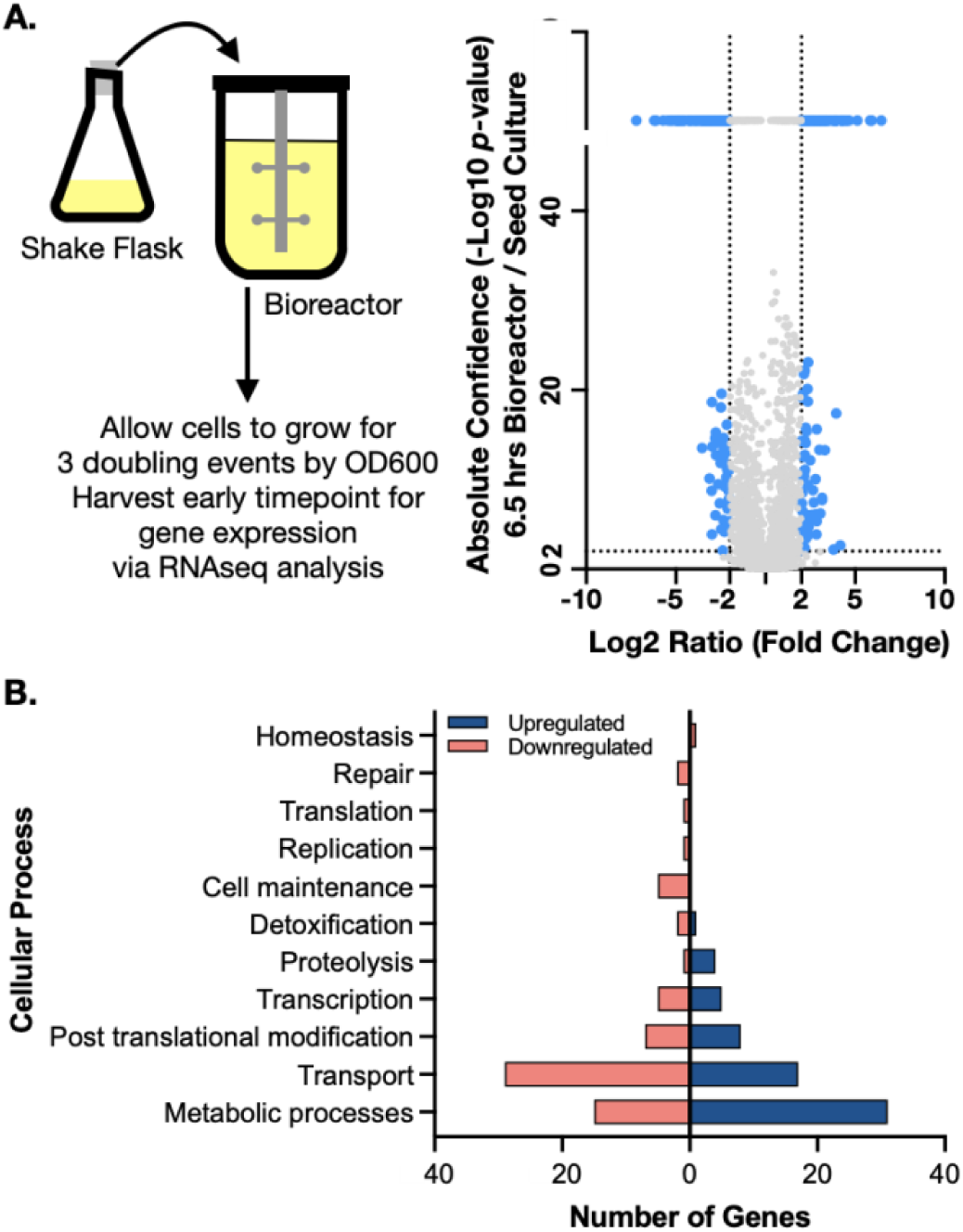
Genome wide expression differences in diverse cellular processes upon shifting to a stirred tank bioreactor. **(A)** *Left side*. Schematic showing scale transition from 25mL seed culture of IP producing *C. glutamicum* in CGXII media to a stirred tank bioreactor. *Right side*. Volcano plot comparing differential gene expression (6.5 h post inoculation / shake flask) via RNAseq analysis to absolute confidence (*p* value) of the same time points. Fold changes greater than 4 (log2=2) and absolute confidence values >2 (*p*<0.001) are considered significant. The threshold for significance is demarcated with dotted lines and the corresponding genes are colored blue. Genes with insignificant differential expression are indicated in grey. Genes with confidence values >40 are placed above the break on the y axis. **(B)** Analysis of gene classes enriched in the scale transition. Differentially expressed genes from a) were binned into functional categories based on COG annotations and putative function by BLAST alignment. Upregulated genes are indicated in dark blue; downregulated genes are indicated in light red.

Many genes encoding metabolic functions were differentially expressed in the transition from shake flask to stirred tank format. We used a fold change cutoff of 4 (log2 > 2) and a *p* value < 0.001 to identify both large and statistically significant changes. Gene ontology (GO) enrichment annotations identified the highest number of DEGs belonging to metabolism and transport processes (**Figure 3B**). The strongest fold changes (16-fold increase or higher) were in metabolism; Cgl2807 (*adhA*, zinc dependent alcohol dehydrogenase), Cgl1396 (acetylglutamate kinase), Cgl2886 and Cgl2887 (two FAD-dependent oxidoreductases) and Cgl3007 (*mez*, malic enzyme). Of these genes, Cgl2807/*adhA* encodes for a Zn-dependent alcohol dehydrogenase that together with Cgl2796 has been reported to maintain redox balance (Zhang et al., 2018). While the cells had been previously adapted in CGXII medium for the seed culture, we observed differentially increased gene expression of several amino acid biosynthesis pathways. Increased gene expression for nearly complete pathways needed for methionine, leucine, and arginine biosynthesis were detected, as well as the gene responsible for glutamate synthesis, *gdh*. Three genes responsible for the conversion of propionate to succinate and pyruvate through the methylcitrate cycle were also upregulated. Upregulated DEGs encoding for myo-inositol metabolism directing flux towards acetyl-CoA and DHAP included Cgl0163/*iolE*, Cgl0161/*iolB*, Cgl0158/*iolC*, Cgl0160/*iolA/msmA*, and Cgl0157/*iolR*. Of the myo-inositol pathway genes, *iolR* was reported to regulate PTS-independent glucose uptake by repressing the expression of glucokinases in *C. glutamicum* (*Zhou et al*., *2015*).The upregulation of myo-inositol catabolic pathways could be attributed to supplemental yeast extract amended to the CGXII medium in the bioreactor. Yeast extract was added to the bioreactors as it was found to improve IP production when *E. coli* was used as the microbial host (Kang et al., 2019). Inositol is found in the yeast extract (>160 mg/g range) for many commercial preparations.

A wide range of regulatory factors and stress responsive genes were also upregulated at the shake flask to bioreactor transition time point. Cgl2988/*malR*, which encodes for a MarR type transcriptional regulator and Cgl3007/*mez* were both highly upregulated. MalR represses expression of the malic enzyme gene, *mez* (Krause et al., 2012) and is a global regulator of stress-responsive cell envelope remodeling in *C. glutamicum* (Hünnefeld et al., 2019). Cgl2996/*ino-1* (myo-inositol-1-phosphate synthase) is the first enzyme in mycothiol biosynthesis and plays a major role in the detoxification of stress-inducing factors, maintaining the redox balance and protection against oxidative stress (Chen et al. 2019). The universal stress response protein Cgl1407/*uspA2* and HSP 60 family chaperonin, Cgl2716/*groEL* were also upregulated.

A similar number of genes were downregulated during the transition from shake flask to bioreactor **(Figure 3B)**. Of the genes uniquely downregulated at 6.5 h, included Cgl1427/*cmk*, cytidyl kinase, Cgl2605/*bioD*, thioredoxin reductase. Cgl1427 has been reported to be crucial for maintaining triphosphate pools (ATP, CTP) under oxygen-limiting environments (Takeno et al., 2013) but it’s downregulation implies these early time points are not oxygen-limited. Several genes involved in transport were also significantly downregulated with a cutoff threshold log2 ratio less than -4. These included ABC transporter ATPase proteins Cgl1351, Cgl1546/*pacL* (cation specific) and Cgl1567 along with Cgl2222, a major facilitator superfamily (MFS) transporter. Downregulated genes Cgl0026-Cgl0029 have been reported to be Zur-binding sites that are involved in zinc homeostasis in *C. glutamicum* (Schröder et al., 2010). Other downregulated transporters included the lysine exporter Cgl1262/*lysE*, exporter systems for branched chain amino acid and methionine (*brnE/brnF*) along with several MFS transporters (Cgl1065, Cgl1076/*pcaK*, Cgl0380, Cgl0381, Cgl2685/*lmrB*) and the ABC type phosphate uptake system (*pstSCAB*). Several other ABC transporter subunits (permease or substrate-binding domain or the ATPase) responsible for transport of iron, calcium, cobalt, cadmium, copper, sn-glycerol-3-phosphate, etc. were also downregulated. Downregulated transcriptional regulators during this scale transition phase belong to the GntR family (Cgl2316), ArsR family (Cgl2279), PadR family (Cgl2979) and CopY family (Cgl0385). A complete list of DEGs can be found in **Supplementary datasets, Datasets S1** through **S6** and at the JGI Genome Portal (https://genome.jgi.doe.gov/portal/) under Project ID 1203597.

### 2.4. Metabolic Pathway Alterations during Fed-batch Cultivation indicated by differentially expressed genes

After inoculation into the bioreactors, we benchmarked the bioreactor run with online and offline measurements including growth, glucose consumption, and organic acid secretion, with and without [Ch][Lys]. We noted several differences between cells grown in the control reactor and the [Ch][Lys] treated reactor. While cells were pulse-fed the same feed solution to restore glucose levels back to 60 g/L, the [Ch][Lys] treated engineered strain much less acetate and succinate than the control (**Figures 4A** and **5A**). Overall OD_600_ measurements indicated similar initial growth patterns before the first feeding, but after feeding, OD_600_ measurements did not appreciably increase further and instead we detected overflow metabolite accumulation above 10 g/L of succinate and acetate **(Figure 4A)**. The control reactor decreased in OD_600_ from a high of 49 to a 21 OD_600_. The [Ch][Lys] reactor also decreased in OD_600_, but from a similar high of 50 to 36 OD_600_ (**Supplementary Figure S2**). We correlated gene expression changes during this campaign for both reactors using RNAseq analysis to understand how glucose was redirected from growth to the generation of these overflow metabolites (**Supplementary Data, Dataset S2)**.

**Figure 4:**
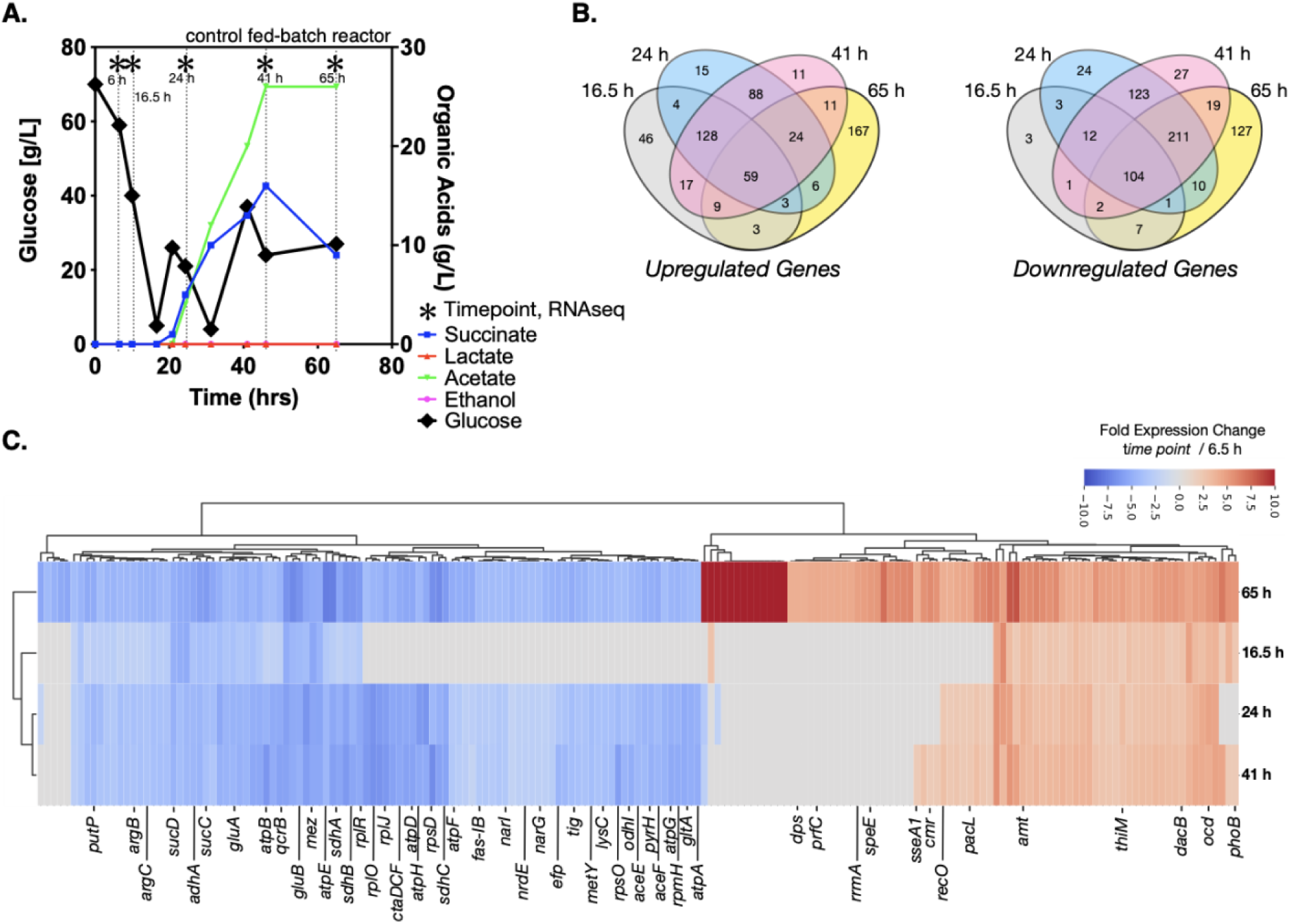
Growth of engineered *C. glutamicum* for isopentenol in a control stir tank bioreactor. **(A)** HPLC analysis of glucose and organic acids detected in the 2L stirred tank bioreactor. Cells were harvested from the indicated time points with (*). Refer to Figure 5a for the [Ch][Lys] treated bioreactor. **(B)** Shared and unique differentially expressed genes. Venn diagrams indicate the number of Upregulated (*left)* and downregulated (*right)* genes at the indicated time points. **(C)** Hierarchical cluster analysis of the top 181 differentially expressed genes at the 65 hr time point vs the 6.5 hr time point for both up or down regulation. A number of genes that are highly expressed only in stationary phase *vs* constitutively expressed are observed.

**Figure 5:**
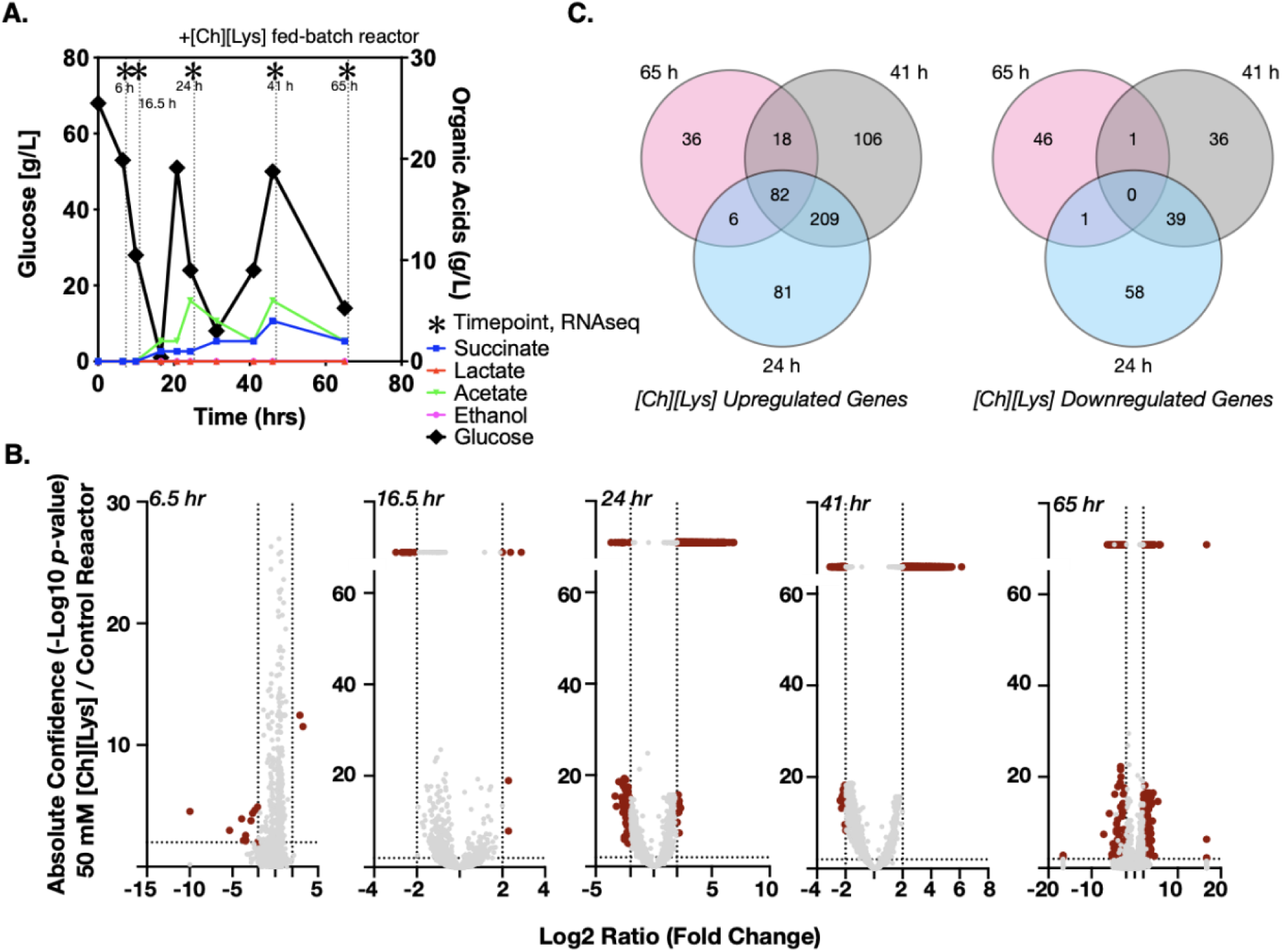
Differential expression of genes in response to 50 mM of [Ch][Lys]. **(A)** HPLC analysis of glucose and organic acids detected in the 2-L stirred tank bioreactor of cells grown in the presence of an initial concentration of 50 mM [Ch][Lys]. Cells were harvested from the indicated time points with (*). Refer to Figure 4a for the control bioreactor. The glucose and organic acid values for the time course in this figure panel have been previously described in Eng and Sasaki *et al*, 2020. **(B)** Volcano plots of differentially expressed genes for each time point. Genes which have confidence values or log2 ratios greater than the maximum value on each axis are plotted on a discontinuous portion of the axis as indicated with a line break. c) Shared and unique differentially expressed genes in response to [Ch][Lys]. Very few differences were detected in the 6 h and 16.5 h time points and are not included in the Venn diagram. DEG was calculated as the ratio between the treated reactor and its corresponding time-matched sample in the other control reactor. Venn diagrams indicate the number of upregulated (*left)* and downregulated (*right)* genes at the indicated time points.

We observed several genes encoding metabolic processes related to succinate and acetate metabolism were downregulated in the time course, such as *ptaA, ackA* and *sucC*. Decreasing their gene expression suggests a decrease in activity, enabling greater succinate or acetate accumulation due to fewer competing reactions for these metabolites as precursors. Cgl2211, a putative succinate exporter (Huhn et al., 2011; Litsanov et al., 2012; Prell et al., 2020) was upregulated at 65 h, that might explain higher succinate excretion profile for the fed-batch cultivation in the absence of the IL (**Figure 4A**). The higher acetate secretion in this bioreactor correlated with upregulated Cgl2066 transcripts at 24 h and 41 h, which encodes a putative acyl phosphatase that converts acetyl phosphate to acetate. At the last phase of cultivation Cgl2380/*mdh* was upregulated (log2 ratio of 3.14) with 12-fold over expression. Malate dehydrogenase, *mdh*, is involved in a NADH based reversible reaction in TCA and is responsible for NADH balance maintenance and succinate formation. The malic enzyme, Cgl3007/*mez*, was downregulated across all later time points (Log2 -3.1 to -7.65), with 10-fold decrease in expression in the last time point alone. Malic enzyme, upregulated during transition from shake flask to a bioreactor scale (log2 ratio of 5.11 at 6.5 h, Section 2.3), is involved in gluconeogenesis important for NADPH regeneration for anabolic processes and pyruvate flux at the cost of carbon loss as one mole of CO_2_. Genes encoding cell division proteins including *mraZ, ftsX, ftsW, ftsE, sepF*, were downregulated for later stage cultivation time points (24 h and later) correlating with the lack of increased OD_600_ after glucose was fed at the 24 hour time point. Cgl1502, a putative MFS transporter (PTS based sugar importer) was upregulated in all later bioreactor cultivation time points. These later time points had many shared downregulated genes, indicating a phenotyping similarity **(Figure 4B)**.

A more comprehensive analysis of differential gene expression indicated that many transporters were upregulated in these bioreactor time points (**Figure 4C**, red colored bars). These included ABC transporters for phosphonate (*pctABCD*); sn-glycerol-3-phosphate (*ugpABCE*) and phosphate (*pstSCAB)*, a branched chain amino acid and methionine exporter (Cgl0258/*brnF*); Cgl0968/*lysI*, which encodes a protein involved in lysine uptake (Seep-Feldhaus et al., 1991). Transcriptional regulators that were upregulated across all the later time points of the bioreactor cultivation and were associated with putative functions included Cgl2496/PucR family, Cgl0962/TetR family, Cgl2934/MarR family, Cgl1367/LacI family and Cgl2616/LysR family. Cgl2776 which is a putative XRE family transcriptional regulator MsrR was found to be upregulated from 24 h to 65 h. *msrR* is located downstream of the *cmr* gene that encodes for a MFS multidrug efflux protein and upstream of Cgl2775/*sseA1*, a sulfurtransferase and Cgl2774. These late-phase upregulated genes have been previously reported to be regulated by MsrR and overexpressed in response to oxidative stress response in *C. glutamicum* (Si et al., 2020). Genes under the control of DtxR, a master regulator of iron homeostasis at late exponential phase (Küberl et al., 2020), and AmtR, a master regulator of nitrogen metabolism (Beckers et al., 2005) were also upregulated at later time points compared to 6.5 h. The iron homeostasis genes included Cgl0387 (putative membrane protein) and Cgl2035, an ABC-type cobalamin/Fe3+-siderophores transporter. The nitrogen metabolism regulon included genes encoding for ammonium permease, *amt*; a predicted ornithine decarboxylase (*ocd*) and the ABC transporter for urea UrtABCDE. Ammonium is a critical precursor for growth and tetramethylpyrazine (TMP) production (Xiao et al., 2014).

We also observed significant downregulation of *adhA, ald, sucCD, malE/mez* **(Figure 4C**, blue colored genes), which were previously reported during microaerobic aeration in a bioreactor cultivation of *C. glutamicum* (Lange et al., 2018). A different complement of transporter-related genes was also downregulated across all the later time points that included genes encoding for maltose and trehalose ABC transporter subunits (Cgl2460 and Cgl0727) and the entire glutamate ABC transporter operon *gluABCD*. This expression profile suggests that at the cell density reached by 20 hours, there was a general cell stress response and the activation of microaerobic-specific genes. The growth conditions did not promote additional cell growth due to the downregulation of cell division genes; glucose uptake genes were still highly active, enabling a significant conversion of glucose to organic acids but not biomass accumulation.

We observed a unique class of genes that were only expressed after high accumulation of succinate and acetate at the 65 hour time point. At this time point, glucose consumption has stalled, and the overflow organic acids have plateaued at the ∼10 g/L concentration. Genes encoding for ROS detoxification including catalase gene Cgl0255/*katA*, superoxide dismutase gene Cgl2927/*sod* along with Cgl2003/*gor*, a mycothione reductase involved in arsenate detoxification were upregulated. DEGs that were downregulated included genes encoding for *catA2, catC, nagI, qsuB, benC and benD*. These are enzymes involved in aromatic compound degradation through beta ketoadipate pathway that would reroute flux into TCA through succinate and acetyl CoA. We interpret the expression of these genes as indicative of the unfavorable cell growth conditions.

A regulator involved in diverting acetyl CoA flux towards fatty acid biosynthesis, Cgl2490/*fasR* was constitutively expressed up until the last time point during bioreactor cultivation in absence of IL. This TetR type transcriptional regulator controls fatty acid biosynthesis and malonyl CoA formation from acetyl CoA and has been deleted for improving malonyl CoA production (Milke et al., 2019). Our analysis correlated this repression by *fasR* with down regulated Cgl2495/*fas-IA* as well as downregulation of Cgl0700/*accBC*, Cgl0708/*dtsR1* and Cgl0707/*dtsR2* during later time points in absence of IL.

### 2.5. C. glutamicum exhibits a complex response to the IL, Cholinium Lysinate under fed-batch cultivation in the bioreactor

Next we analyzed differential gene expression when cells were grown in the presence of 50 mM [Ch][Lys], simulating hydrolysate prepared under a water-conservation regimen (Neupane et al., 2017). Ionic liquids have been reported to increase osmotic pressure, attack lipid structures and consequently disrupt microbial membranes (Pham et al., 2010; Khudyakov et al., 2012; Yu et al., 2016). *C. glutamicum* exhibited differential expression of 727 genes (**[Supplementary Data](https://drive.google.com/file/d/1w_ThbEcxdX8Auizt2bYSxTJl6scE69L8/view?usp=sharing), Dataset S3**), during the [Ch][Lys] treated fed-batch bioreactor cultivation in comparison to the untreated culture at the time-matched samples (**Figure 5A**). While both bioreactors consumed the initial glucose in the reactor at similar rates, their response to the first feeding at 24 hours differed. The [Ch][Lys] reactor showed maximum accumulation of 4 g/L succinate and 6 g/L acetate over the duration of this time course, a 4 fold decrease for both organic acids in the absence of [Ch][Lys] (compare **Figure 4A** to **Figure 5A**). In the presence of [Ch][Lys], genes encoding for succinate utilization such as *sdhA, sdhB* and *sdhC* were all upregulated at 24 h and 41 h in contrast to the control reactor. Similarly, genes encoding for pyruvate decarboxylation to acetyl CoA (instead of acetate) via *aceE* and *aceF* were also highly upregulated at later time points.

During the early cultivation time points (6.5 - 16.5 h), only 1.5% of the total pool of differentially expressed genes changed in response specifically to [Ch][Lys], but the datasets diverged after the first feeding at 24 hours as biomass formation reached its maximum (**Figure 5B**). Only two genes were upregulated at the 6.5 h time point: a MFS transporter (Cgl2611) and its transcriptional regulator (Cgl2612) (**Supplementary Data, Dataset S3**). The BRC-JBEI 1.1.2 homolog is 97.37% identical to Cgl2611 which exports cadaverine, a L-lysine derived product (Kind et al., 2011; Adkins et al., 2012; Jones et al., 2015; Tsuge et al., 2016). Cgl2611 expression was not detected at the control 6.5 h time point, but both genes are highly upregulated with or without [Ch][Lys] treatment in the remaining time points. Cgl1203, which encodes a phospho-N-acetylmuramoyl-pentapeptide-transferase associated with cell wall biosynthesis, was only upregulated at 16.5 h.

Early transcriptome changes in *C. glutamicum* during bioreactor cultivation post [Chl][Lys] exposure included overexpression of MFS transporters along with repression of mechanosensitive channels that were consistent with IL tolerance mechanisms reported in other microbes (Khudyakov et al., 2012; Martins et al., 2013; Yu et al., 2016). Many genes were downregulated in response to exogenous [Ch][Lys] in the bioreactor and represented 25% of DEGs. Cgl0879/*mscL*, a large-conductance mechanosensitive channel, and is related to osmotic regulation (Krämer, 2009), was uniquely downregulated at 16.5 h.

A comprehensive analysis of upregulated DEGs at more than one time point represented around 59% of the total upregulated genes in the presence of IL (**Figure 5B**). Nearly 15% of those genes showed consistent overexpression from 24 h through 65 h (**Figure 5C)**. This differential transcript profile reflects the metabolic perturbation over the course of the fed-batch cultivation after the initial glucose exhaustion followed by glucose pulse feeding and is depicted in **Figure 6**. Prominent DEGs include those encoding for energy metabolism, amino acids biosynthesis, response to oxidative and other environmental stress conditions (**Figure 5E** and **Supplementary Data, Dataset S4**). Genes involved in energy metabolism were highly upregulated during the later phase of fed-batch cultivation in the presence of IL compared to its absence. These included NADH dehydrogenase (Cgl1465), succinate dehydrogenase, *sdhABC* genes at 24 h and 41 h; cytochrome oxidase, *ctaDCEF*, cytochrome reductase, *qcrCAB* and the ATP synthase complex (Cgl1206 to Cgl1213) genes at 24 h, 41 h and 65 h. Amino acid biosynthetic genes upregulated at the later time points included the arginine biosynthetic genes *argC argJ, argB* and *argH* at 65 h and *argG* and *argD* at mid cultivation phase (41 h). ArgJ protein was also enriched in the acetoin/TMP producing *C. glutamicum* strain (Eng et al., 2020). Genes encoding for other amino acid biosynthesis included Cgl1139/*metE*, Cgl2446/*metB* and Cgl0653/*metY* at 24 h and 41 h from the methionine/homocysteine pathway; Cgl2204/*ilvE* at 24 h and Cgl1273/*ilvC* at 24 h and 41 h in the branched amino acid pathway. Several ribosomal proteins were significantly upregulated during the same cultivation phase (24 h and 41 h) including 30S ribosomal proteins S15 (Cgl1976/*rpsO*) and S18 (Cgl0866/*rpsR*); 50S ribosomal proteins L28 (Cgl0869/*rpmB*) and L15 (Cgl0542/*rplO*) along with the ribosome recycling factor Cgl2023/*frr*.

**Figure 6:**
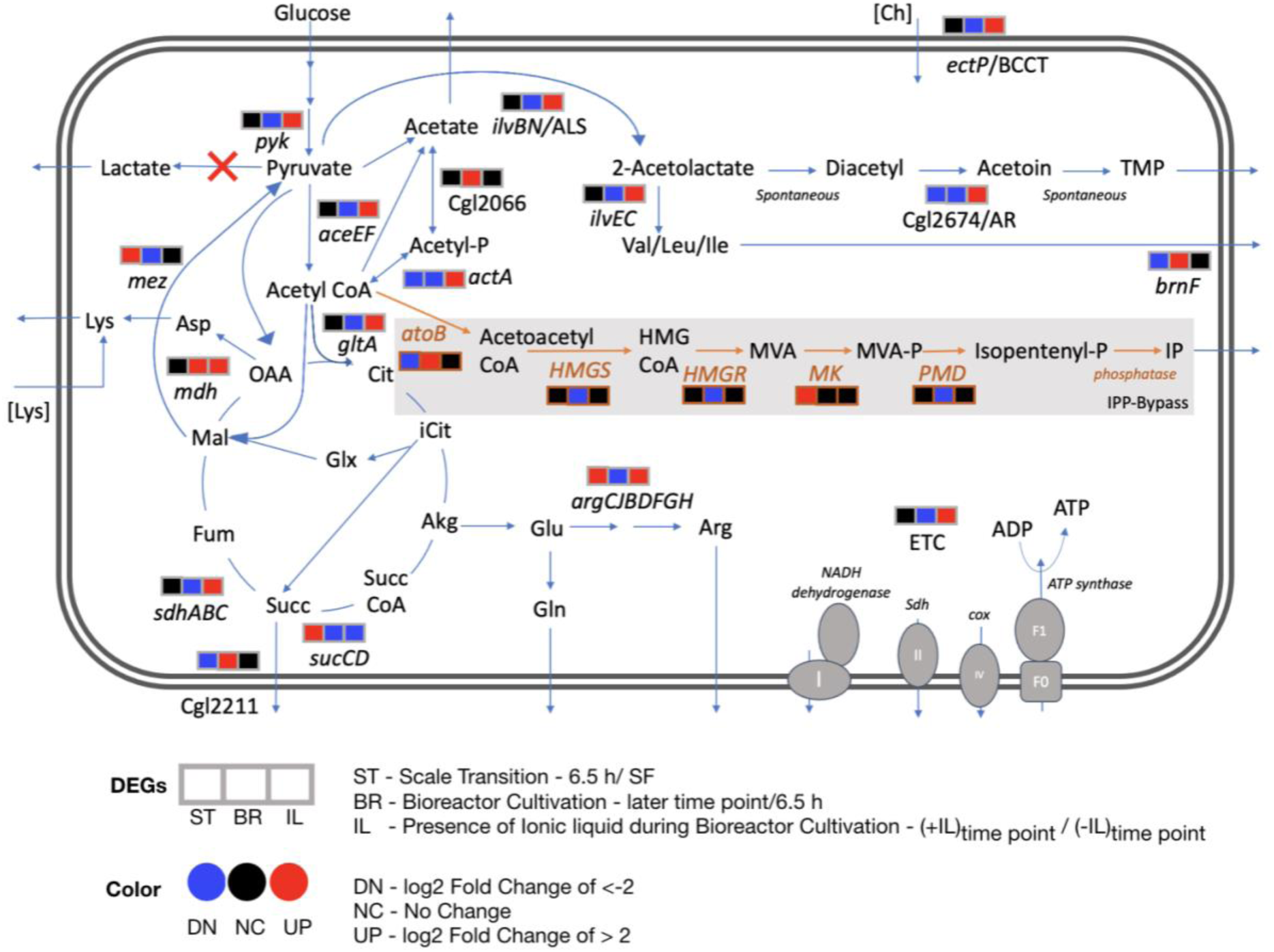
Differential transcript profiles of engineered *C. glutamicum* under fed-batch cultivation. Three DEGs corresponding to three discrete conditions that were analyzed are represented here: ST - scale transition from shake flask (SF) to early bioreactor cultivation (6.5 h), BR - bioreactor later stage cultivation in the absence of IL and IL - bioreactor cultivation in the presence of IL compared to in the absence of IL. The heterologous pathway for IP production is shown in orange. Red crosses show the gene deletions in the *C. glutamicum* strain used in this study. Abbreviations: Acetyl-P, acetyl phosphate; Akg, alpha ketoglutarate; Arg, arginine; Asp, aspartate; *atoB*, acetyl-CoA acetyltransferase; Cit, citrate; Ch, cholinium; Cox, cytochrome oxidase; ETC, Electron transport chain; Fum, fumarate; Glx, glyoxylate; Glu, glutamate; Gln, glutamine; *HMGS*, hydroxymethylglutaryl-CoA synthase; *HMGR*, 3-hydroxy-3-methylglutaryl-CoA reductase; HMG CoA, 3-hydroxy-3-methyl-glutaryl-coenzyme A; Icit, isocitrate; IP, Isopentenol; Lys, lysine; Mal, malate; *MK*, mevalonate kinase; MVA, mevalonate; MVA-P, mevalonate; OAA, oxaloacetate; PMD, phosphomevalonate decarboxylase; Succ, succinate; Succ CoA, succinyl-CoA; Sdh, succinate dehydrogenase; TMP, tetramethylpyrazine.

We also observed the upregulation of an ABC transporter (Cgl0946 and Cgl0947), a multidrug transport system (MTS) operon, in part regulated by its adjacent two-component system (TCS) (Cgl0948-Cgl0949, also upregulated). MTS offers a natural defense against toxic compounds and is reported to be upregulated in response to the non-ionic surfactant Tween 40 (Jiang et al., 2020). Also, Cgl2312/*ectP*, a putative BCCT family transporter was overexpressed in the bioreactor with IL at 24 h time point. This gene, an orthologue for *betT* gene in *E. coli* and *P. putida*, was under-expressed in the bioreactor without IL at later time points (24 h, 41 h). Betaine/carnitine/choline (BCCT) family transporters could enable cholinium uptake and catabolism. An array of other transporters and transcriptional regulators were also downregulated in the presence of IL (**Supplementary Data, Dataset S3**).

While the analysis above compared matched time points with or without [Ch][Lys] treatment, we also included one additional analysis to examine DEGs from samples in the same reactor but as they progressed from the 41 h to 65 h time point (**Supplementary Figure S3**, and **Supplementary Data, Dataset S6**). As observed from our earlier analysis in **Figure 4C** a set of DEGs in the control bioreactor were detected, consistent with entry into the stationary phase. Significantly downregulated genes also included genes encoding for a stationary phase repressor protein/redox responsive transcription factor, *whiB/*Cgl0599 (Walter et al., 2020) and a branched chain amino acid transporter (Cgl2250) (Graf et al., 2019). Cgl2250 has been reported to be downregulated during the transition from exponential to stationary phase in *C. glutamicum* (Larisch et al., 2007).

### 2.6. Indication of Flux rerouting in the presence of IL stress during fed-batch bioreactor cultivation

Our transcriptome analysis identified differential profiles for energy metabolism, amino acid biosynthesis and redox related genes as discussed in the previous section (**Figure 6**). Several genes encoding for metabolic reactions related to acetoin and TMP accumulation were specifically upregulated in the presence of 50 mM of [Ch][Lys] at the 24 h or 41 h time points (**Supplementary Data, Dataset S3, Supplementary Figure S4**) when compared to the control samples at the same time points. Of the two subunits of the acetolactate synthase (ALS) *ilvB and ilvN*, the smaller regulatory subunit, Cgl1272/*ilvN* was upregulated in the presence of IL fed-batch cultivation when compared to the absence of IL at 24 h. Acetolactate synthase in *C. glutamicum* takes part in diverting pyruvate flux towards branched chain amino acids biosynthesis and acetoin biosynthesis and could be a precursor to TMP (Eng et al., 2020) **(Figure 5**). Although branched chain amino acid biosynthesis has been extensively researched for engineering branched chain alcohol (e.g. isobutanol) producing *C. glutamicum* strains (Hasegawa et al., 2020) the branched chain amino acid degradation towards isopentenol biosynthesis (through HMG-CoA) and TCA through acetyl CoA still remains to be fully investigated. The other proposed enzyme in TMP accumulation is the NADH consuming acetoin reductase (AR, Cgl2674) and was also significantly upregulated (log2>4) at 41 h in presence of 50 mM of [Ch][Lys] compared to fed-batch cultivation in the absence of IL at similar time points. Genes encoding mechanisms that divert pyruvate flux towards acetyl CoA (Cgl2248/*aceE* and Cgl2207/*aceF*) were also upregulated along with genes for pyruvate kinase (Cgl2089/*pyk*) and citrate synthase (Cgl0829/*gltA*).

## 3. Discussion

*C. glutamicum* is a strong contender as a microbial chassis for IP production and is already used at commercial scales. To test IP production in stirred-tank bioreactors, we used process optimizations empirically identified for high IP titers in *E. coli* (Kang et al., 2019). Using this alternative microbe, Kang et al reported IP titers > 3 g/L in fed-batch mode production; in contrast, these process parameters led to much lower IP titers in our *C. glutamicum* strains and were instead near the lower detection limit. It is possible that these optimizations were specific to *E. coli;* the impact of this IP production pathway in *C. glutamicum* upon shifting from batch mode to fed-batch mode in a stirred tank bioreactor may have resulted in a different host-specific metabolic response.

What parameters are important in selecting one microbial host over another? From a genetic tractability perspective, the biggest drawback of *C. glutamicum vs*. model microbes such as *E. coli* could arise from its reduced transformation efficiency, which was lower by 3-5 orders of magnitude (Chung et al., 1989; Inoue et al., 1990; Ruan et al., 2015). However, Baumgart and coworkers made an astute observation; by using a methylation deficient strain of *C. glutamicum*, one could both improve transformation efficiency as well as plasmid copy number (Baumgart et al., 2013). Improved pathway copy number (both genomically integrated or plasmid-borne) in *E. coli* had already been shown to dramatically improve heterologous isoprenoid titers (Goyal et al., 2018; Chatzivasileiou et al., 2019). With this premise we initially used a methylation deficient strain as our starting host. However, the methylation deficient strain only produced trace titers of IP but a related strain produced both improved IP titers 20x or a co-product, tetra-methylpyrazine. Understanding the genetic differences in this isolate BRC-JBEI 1.1.2 was the major thrust of this study.

Leveraging strain isolate differences is already commonplace when analyzing natively expressed products, such as natural products from *Streptomyces* spp. or wine, beer, and baking in *Saccharomyces* spp. (Nepal and Wang, 2019; Gallone et al., 2016). In *E. coli*, the Hanahan cloning strain DH1 is the preferred strain for the production of many terpenes, but experimentally identified modifications are needed to translate port pathways to other *E. coli* isolates as with the case for limonene production in *E. coli* BL21(DE3) (Tsuruta et al., 2009; Rolf et al., 2020). A potential explanation for DH1 being a more robust host may be due to its elevated number of ribosomes compared to strains DH10, BL21, or BW25113 (Cardinale et al., 2013), which may indirectly help with heterologous pathway protein expression. Our whole-genome sequencing analysis identified a large number of genetic differences in our engineered isopentenol producing *C. glutamicum* BRC-JBEI 1.1.2 isolate (many associated with metabolic functions) that are unaccounted for when using the reference *C. glutamicum* genome. Previously we used computationally driven maximum theoretical yields calculations for a product across several microbes to evaluate microbial potential for a specific product/substrate pair (Banerjee et al., 2020). However the accuracy of such predictions rely on the metabolic reactions curated for the reference strain, and are challenging to apply in isolates used with differences at the genomic or metabolic level (refer to IP titers in **Figure 1A**). Pan-genome assemblies and metabolic models can be applied to this situation (both for BRC-JBEI 1.1.2 and DH1) to more accurately account for these metabolic features (Monk et al., 2013; Norsigian et al., 2018).

For emerging processes using IL pretreated lignocellulosic biomass, *C. glutamicum* as the microbial IP producer for this process is compelling. To the best of our knowledge, this is the first transcriptomics analysis of an engineered isopentenol producing *C. glutamicum* strain in fed-batch conditions. Due to the relative similarity between this isolate to the type strain, we were able to use existing gene annotations with a fairly low homology cutoff (>70%) for the majority of detected transcripts in this study. A large number of significant DEGs identified in this analysis encode hypothetical proteins that lack functional information. These genes can be further characterized using functional genomics tools such as parallelized transposon mutant libraries (Lim et al., 2019; Cain et al., 2020) or high throughput transcription factor characterization (Rajeev et al., 2014, 2011) to improve our understanding of these useful *C. glutamicum* isolates. Transcriptomics analysis completed here indicated that in order to improve isopentenol titers under stirred tank fed-batch conditions, targeting deleting *mdh* could limit accumulation of succinate, a highly overexpressed gene. *gltA*, Cgl2211, *brnF* and arginine biosynthesis genes were also highly upregulated (**Figure 6**); deleting or down regulating them could enlarge the acetyl-CoA pool, in turn improving IP titers. Additional gene targets include *pta-acka, poxB, actA* and Cgl2066 to block acetate formation. This transcriptomics analysis also implicated *ectP*, a BCCT family transporter similar to *E. coli betT* and *P. putida betT-III*, as a transporter for [Ch][Lys]; *ectP* was overexpressed in the presence of ILs. A BCCT transporter has been proposed to be involved in uptake and catabolism of the cholinium ion from [Ch][Lys] in both *E. coli* and *P. putida* (Park et al., 2020). Characterizing IL tolerance is an active research thrust in our laboratory.

In summary, our transcriptomic analysis under industrially relevant process conditions provides a toehold for future DBTL cycles. Future “learn” steps can leverage the information gleaned here to target the critical features implicated for improved *C. glutamicum* strain performance when producing desirable products, like isopentenol. Even accounting for potential increased cell heterogeneity in the bioreactor (Wehrs et al., 2019), important features both common and unique to conditions allow a closer look into cell physiology.

## 4. Materials and Methods

### 4.1. Reagents and Experimental conditions

In a previous report (Sasaki et al., 2019), we referred to the IP producing *C. glutamicum* strain as ATCC 13032 NHRI 1.1.2, as indicated in our archival notes. As we cannot confirm the provenance of *C. glutamicum* BRC-JBEI 1.1.2 and how it may have been derived from its closest relatives *C. glutamicum* SCgG1 or SCgG2, we opted to give this strain a unique identifier to avoid further confusion.

Unless indicated elsewhere, all reagents used were molecular biology grade or higher. Primers were synthesized by IDT DNA Technologies (Coralville, IA). CGXII media was prepared as previously described (Sasaki et al., 2019; Keilhauer et al., 1993). All strains and plasmids used in this study are described in **Supplementary Table S2**. *C. glutamicum* strains were struck to single colonies from glycerol stock on LB plates containing the appropriate antibiotic and prepared for production runs as previously described (Eng et al., 2020). The fed-batch cultivation with 50 mM of [Ch][Lys] supplementation was previously described in (Eng et al., 2020). The control bioreactor without [Ch][Lys] was conducted at the same time and the glucose feeding regime was identical to that of the ionic liquid (IL) supplemented reactor. For RNAseq extraction, 5mL culture samples were harvested in 1 mL aliquots, collected by centrifugation at 14,000x*g* for 3 minutes, and stored at -80 °C until subsequent RNA extraction. The supernatant from one of the appropriate time point aliquots was processed for organic acid analysis as described previously (Eng et al., 2020). Lab-scale IP production runs in deep well plates or 5 mL culture tubes were conducted as previously described (Eng et al., 2020). Isopentenol titers reported for the deep well plate format were corrected for evaporation at the 48 h time point as conducted previously (Sasaki et al., 2019). Exogenous [Ch][Lys] toxicity against *C. glutamicum* ATCC13032 and BRC-JBEI 1.1.2 was analyzed in a 48-well microtiter dish format. Cells were first adapted two times in CGXII minimal media with 4% (w/v) D-glucose. When cells were back diluted into fresh media in the microtiter dish, the starting OD600 was set to 0.1 with a fill volume of 200 μL. The plate was incubated with shaking at 30 °C and exogenous [Ch][Lys] added at the start of the time course. OD was monitored at 600 nm on a Synergy 4 plate reader (BioTek Instruments, Winooski VT) with the continuous shaking setting.

### 4.2. Production run with ensiled sorghum hydrolysate

CGXII minimal media was supplemented with ensiled sorghum biomass hydrolysate to test the ability of *C. glutamicum* BRC-JBEI 1.1.2 to utilize carbon sources from renewable feedstock pretreated with IL. Briefly, the forage sorghum (NK300 type, grown in Fresno, CA) was planted in Spring 2020 and harvested in Fall 2020. A forage harvester was used to both harvest and chop the sorghum biomass, which was then loaded in a silage pit, inoculated, and covered to maintain anaerobic conditions. The pit was opened in November 2020 and a sample of the ensiled material was collected, packed with dry ice while in transit, and stored at 4 °C. A 210 L scale Andritz Hastelloy C276 pressure reactor (AG, Graz, Austria) with a helical impeller was utilized to process ensiled sorghum for the pretreatment and saccharification processes. Ensiled sorghum biomass was pretreated at 20% w/w solid loading with 10% w/w [Ch][Lys] at 140 °C for 3 h with a mixing speed of 30 rpm. Solid loading was calculated based on the dry matter content determined using a Binder VDL115 vacuum oven. After 3 hours at the target temperature, the reactor was cooled to room temperature before proceeding with the next steps. The Andritz reactor is sealed during this process, preventing contamination until further processing. Following pretreatment, the pretreated materials were adjusted to pH 5.1 using 50% v/v sulfuric acid and an enzyme cocktail of Novozyme, Inc Cellic Ctec3 and Cellic Htec3 commercial enzymes in a ratio of 9:1 was added. Concentration of the commercial stocks were determined using Bradford assays and bovine serum albumin as a reference. Enzyme load was conducted at a ratio of 10 mg enzyme per 1 g of dry weight biomass. Following pH adjustment and enzyme addition, RODI water was added to obtain a final solid loading of 18.70%. Saccharification by enzymatic hydrolysis was operated at 50 °C, 30 rpm for 70 h (Barcelos et al., 2021). The hydrolysate was then sequentially filtered using a filter press through 5 μm, 1 μm, and 0.25 μm filters. Final filter sterilization was completed with a 0.2 μm filter and stored at -80 °C until further use. This hydrolysate was thawed and added in place of water in CGXII media (amounting to 2.8 % (w/v) glucose), pH was adjusted to 7.4 and filter sterilized one additional time before use. We make the assumption the hydrolysate contained no biologically available nitrogen. To maintain a C/N ratio of glucose/ammonium sulfate + urea of 2.8, pure glucose powder was supplemented to the hydrolysate CGXII cultivation medium composition (Sasaki et al., 2019).

### 4.3. DNA and RNA Isolation

Genomic DNA from *C. glutamicum* BRC-JBEI 1.1.2 was isolated with the following protocol. In brief, strains from glycerol stocks were struck to single colonies on LB plates grown at 30 °C overnight. A single colony was then inoculated into a 250 mL shake flask with 25 mL LB media and grown overnight to saturation. Cells were collected by centrifugation at 4,000x*g* for 5 minutes. The cell pellet was then resuspended in 2 mL lysis buffer (2mM EDTA, 250mM NaCl, 2% (w/v) SDS, 2% (v/v) Triton-X 100, 2% (v/v) Tween-80, 5 mM DTT, 30 units Zymolyase 100T, 1 mg/mL RNaseA). Zymolyase was supplied by US Biological (Salem, MA). The cells were initially incubated at 50 °C to promote protease activity and then incubated for an additional 3 hours at 37 °C with occasional mixing, at which point the lysate became noticeably viscous. DNA was extracted following standard protocols for isolation of DNA using phenol chloroform: isoamyl alcohol and subsequent isopropanol precipitation (Sambrook and Russell, 2001).

RNA was extracted from *C. glutamicum* samples using a Direct-Zol RNA Kit (Zymo Research, Irvine, CA) following the manufacturer’s protocol. *C. glutamicum* cells were lysed after initially resuspending the cell pellet in 500 µL TRI reagent and mixed with glass beads. This mixture was then subject to cell disruption using a bead-beater (Biospec Inc, Bartlesville, OK) with a 3 minute homogenization time at maximum intensity. After bead beating, samples were collected following the manufacturer’s protocol without any additional modifications. RNA quality was assessed using a BioAnalyzer (Agilent Technologies, Santa Clara, CA) before RNA library preparation and downstream analysis.

For 16S ribosomal sequencing, *C. glutamicum* ATCC 13032 Δ*mrr* and *C. glutamicum* JBEI-BRC 1.1.2 were struck from glycerol stocks to single colonies on LB plates and incubated overnight at 30 ° C. A single colony was isolated and boiled in 50 µL dH_2_0 for 10 minutes. 1 µL of the boiled colony was used for PCR with primer pair (JGI_27F: 5’-AGAGTTTGATCCTGGCTCAG-3’and JGI_1391R: 5’-GACGGGCRGTGWGTRCA-3’) with NEB Q5 Polymerase (New England Biolabs, Ipswitch, MA). The PCR amplicon was confirmed by agarose gel electrophoresis and the sequence was determined using conventional Sanger Sequencing (Genewiz LLC, Chelmsford, MA).

### 4.4. PacBio Genome Assembly

DNA sequencing was generated at the DOE Joint Genome Institute (JGI) using the Pacific Biosciences (PacBio) sequencing technology. A Pacbio SMRTbell(tm) library was constructed and sequenced on the PacBio Sequel and PacBio RS II platforms, which generated 397,096 filtered subreads (1,418,602,725 subread bases) totaling 3,352,276 bp. The mean coverage for this genome was 432.21x. All general aspects of library construction and sequencing performed at the JGI can be found at http://www.jgi.doe.gov.

### 4.5. RNAseq Library Generation and Processing for Illumina NGS

Stranded RNAseq library(s) were created and quantified by qPCR. Sequencing was performed using an Illumina instrument (refer to Sample Summary Table for specifics per library). Raw fastq file reads were filtered and trimmed using the JGI QC pipeline resulting in the filtered fastq file (* .filter-RNA.gz files). Using BBDuk (https://sourceforge.net/projects/bbmap/), raw reads were evaluated for artifact sequence by kmer matching (kmer=25), allowing for 1 mismatch and detected artifacts which were trimmed from the 3’ end of the reads. RNA spike-in reads, PhiX reads and reads containing any Ns were removed. Quality trimming was performed using the phred trimming method set at Q6. Following trimming, reads that did not meet the length threshold of at least 50 bases were removed.

Filtered reads from each library were aligned to the reference genome using HISAT2 version 2.2.0 (Kim et al., 2015). Strand-specific coverage bigWig files were generated using deepTools v3.1 (Ramírez et al., 2014). Next, featureCounts (Liao et al., 2014) was used to generate the raw gene counts (counts.txt) file using gff3 annotations. Only primary hits assigned to the reverse strand were included in the raw gene counts (-s 2 -p --primary options). Raw gene counts were used to evaluate the level of correlation between biological replicates using Pearson’s correlation and determine which replicates would be used in the DEG analysis (**Supplementary Figure S5**). In the heatmap view, the libraries were ordered as groups of replicates. The cells containing the correlations between replicates have a purple (or white) border around them. For FPKM and TPM, normalized gene counts refer to SRA reads (Data availability section). A sample legend and description of RNAseq libraries used in this paper is described in **Supplementary Table S3**.

### 4.6. Transcriptome Analysis

Global transcriptome response under various experiment conditions were measured using Geneious Prime 2021 (https://www.geneious.com). The normalized expression was calculated and the differentially expressed genes (DEGs) were filtered for absolute log2 ratio > 2 (i.e. a 4-fold up or down regulation), absolute confidence >3 (p<0.001) and >90% sequence identity. The DEGs at various conditions were functionally annotated using Blast2GO suite (Götz et al., 2008) to assign GO annotations (Galperin et al., 2014). Each DEG was subjected to pathway analysis using the KEGG (Kyoto Encyclopedia of Genes and Genomes) database (http://www.kegg.jp/kegg/pathway.html) to explore the biological implications. Biocyc (https://biocyc.org/) was used to calculate pathway enrichment for the last 65 h/41 h time point and for additional gene orthologs identification. Pathways were considered significant if p<0.05. Hierarchically clustered heat maps were generated with average linkage method and euclidean distance metric in Jupyter notebook using Python library Seaborn 0.11.1 (Waskom et al., 2020).

## 5. Data Availability Statement

All datasets generated in this study are included in the article and Supplementary material. The RNAseq datasets generated and analyzed for this study can be found at the JGI Genome Portal under Project ID 1203597. RNAseq datasets have also been deposited at the NCBI SRA database under the following sample accession numbers: SRP239962; SRP239973; SRP239963; SRP239971; SRP239972 ; SRP239970 ; SRP239968 ; SRP239969 ; SRP239966 ; SRP239967 ; SRP239964 ; SRP239965. The draft genome assembly of *C. glutamicum* BRC-JBEI 1.1.2 has been deposited at the NCBI BioProject database with accession number PRJNA533344 and scaffold assembly accession number GCA_011761195.1. The IMG accession number of this genome assembly on the JGI IMG database is 2821586876.

## 6. Supplementary Material

1. List of Supplementary Figures S1 to S5 and Supplementary Tables S1 to S3.
2. Supplementary Data: Datasets S1 to S6

## 8. Author Contributions

Raised Funds: AM BS. Conceptualization of the project: AM TE. Strain construction, molecular biology, bioreactor sample collection and processing: YS TE RH JT. Analytical Chemistry, IP Production Assays, IL toxicity assays: YS, TE, AS. Interpreted results: YS DB TE AM. Contributed critical reagents: NS AO CS DP TE YS JT BS. RNAseq library generation, data collection, validation: VS, YS, TE. Drafted the manuscript: DB TE AM. All authors read, contributed feedback, and approved the final manuscript for publication.

## 10. Acknowledgements

We thank Andrew Lau for feedback on the figures. We also thank Venkata Ramana Reddy Pidatala and Alex Codik for technical assistance.

