## Supplementary Materials for "Genomics Characterization of an engineered *Corynebacterium glutamicum* in Bioreactor Cultivation under Ionic Liquid Stress"

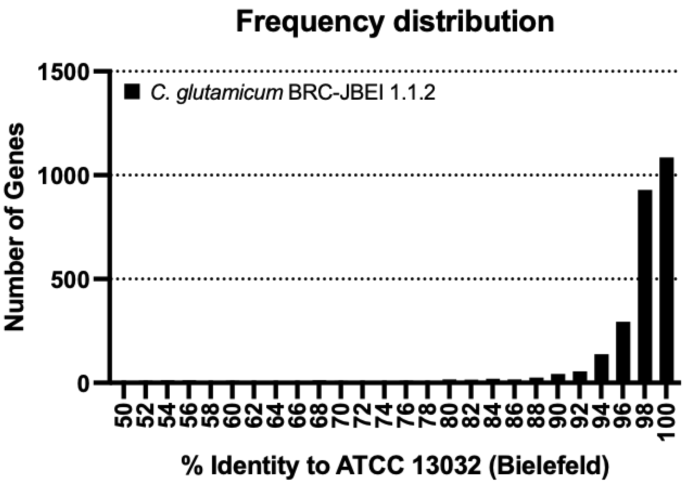


**Figure S1:** Annotating *C. glutamicum* BRC-JBEI 1.1.2 with characterized genes from the C. glutamicum ATCC 13032 (Bielefeld) assembly. 2,777 Cgl genes were annotated in the *C. glutamicum* BRC-JBEI 1.1.2 assembly. Seven genes are in the histogram bin of 50 % identity.

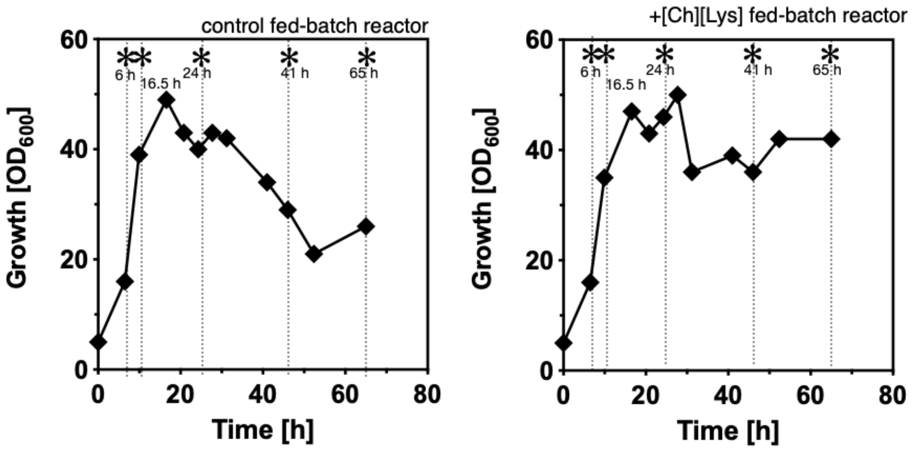


**Figure S2.** Offline $OD_{600}$ measurements of isopentenol producing *C. glutamicum* grown with or without [Ch][Lys] treatment.

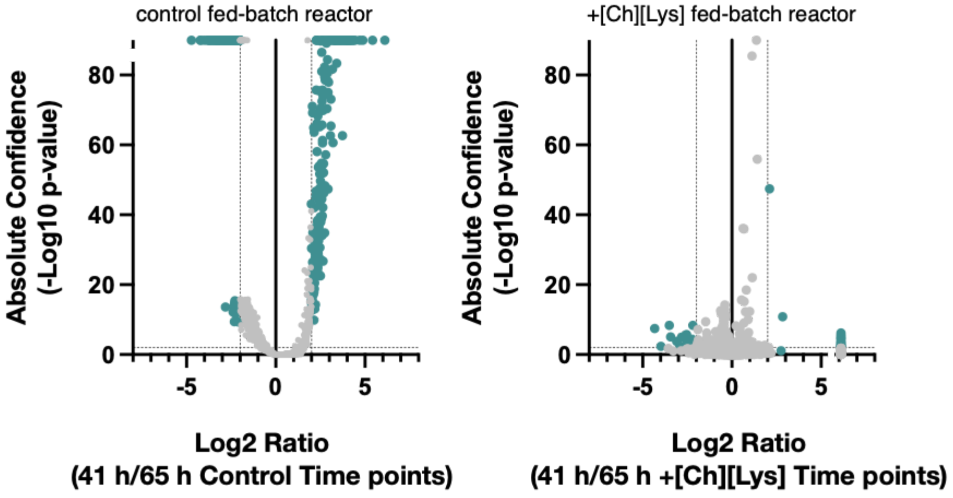


**Figure S3.** Differential gene expression analysis of *C. glutamicum* cells sampled from the same bioreactor nearing termination of fed batch campaign (41 hours / 65 hours) with or without exogenous [Ch][Lys] in culture media.

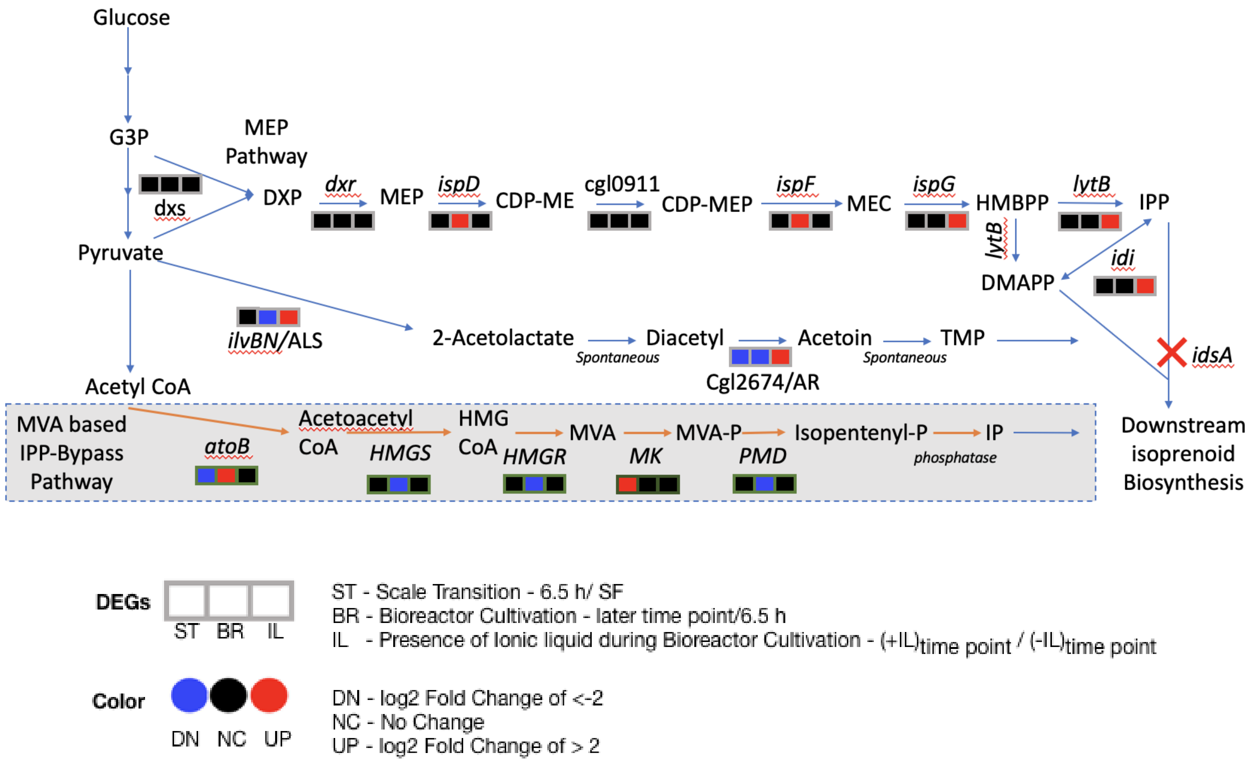


**Figure S4:** Differential transcript profiles of engineered *C. glutamicum* for isoprenoid biosynthesis and acetoin production. The native methylerythritol-4-phosphate (MEP) pathway is compared to the heterologous mevalonate (MVA) based isopentyl diphosphate (IPP) bypass pathway which results in isopentenol (IP) production. Three DEGs corresponding to three discrete conditions that were analyzed are represented here: ST - scale transition from shake flask (SF) to early bioreactor cultivation (6.5 h), BR - bioreactor later stage cultivation in the absence of IL and IL - bioreactor cultivation in the presence of IL compared to in the absence of IL. The heterologous pathway for IP production is shown in orange. Red crosses show the gene deletions in the *C. glutamicum* strain used in this study.

Abbreviations: *atoB*, acetyl-CoA acetyltransferase; CDP-ME, 4-(cytidine 5'-diphospho)-2-C-methyl-D-erythritol; CDP-MEP, 2-phospho-4-(cytidine 5'-diphospho)-2-C-methyl-D-erythritol; DMAPP, Dimethylallyl diphosphate; DXP, 1-deoxy-D-xylulose 5-phosphate; G3P, Glyceraldehyde 3-phosphate; HMBPP, 1-hydroxy-2-methyl-2-(E)-butenyl 4-diphosphate; *HMGS*, hydroxymethylglutaryl-CoA synthase; *HMGR*, 3-hydroxy-3-methylglutaryl-CoA reductase; HMG CoA, 3-hydroxy-3-methyl-glutaryl-coenzyme A; IP, Isopentenol; IPP, Isopentenyl diphosphate; MEC, 2-C-methyl-D-erythritol 2,4-cyclodiphosphate; MEP, 2-C-methyl-D-erythritol 4-phosphate, *MK*, mevalonate kinase; MVA, mevalonate; MVA-P, mevalonate; PMD, phosphomevalonate decarboxylase; TMP, tetramethylpyrazine.

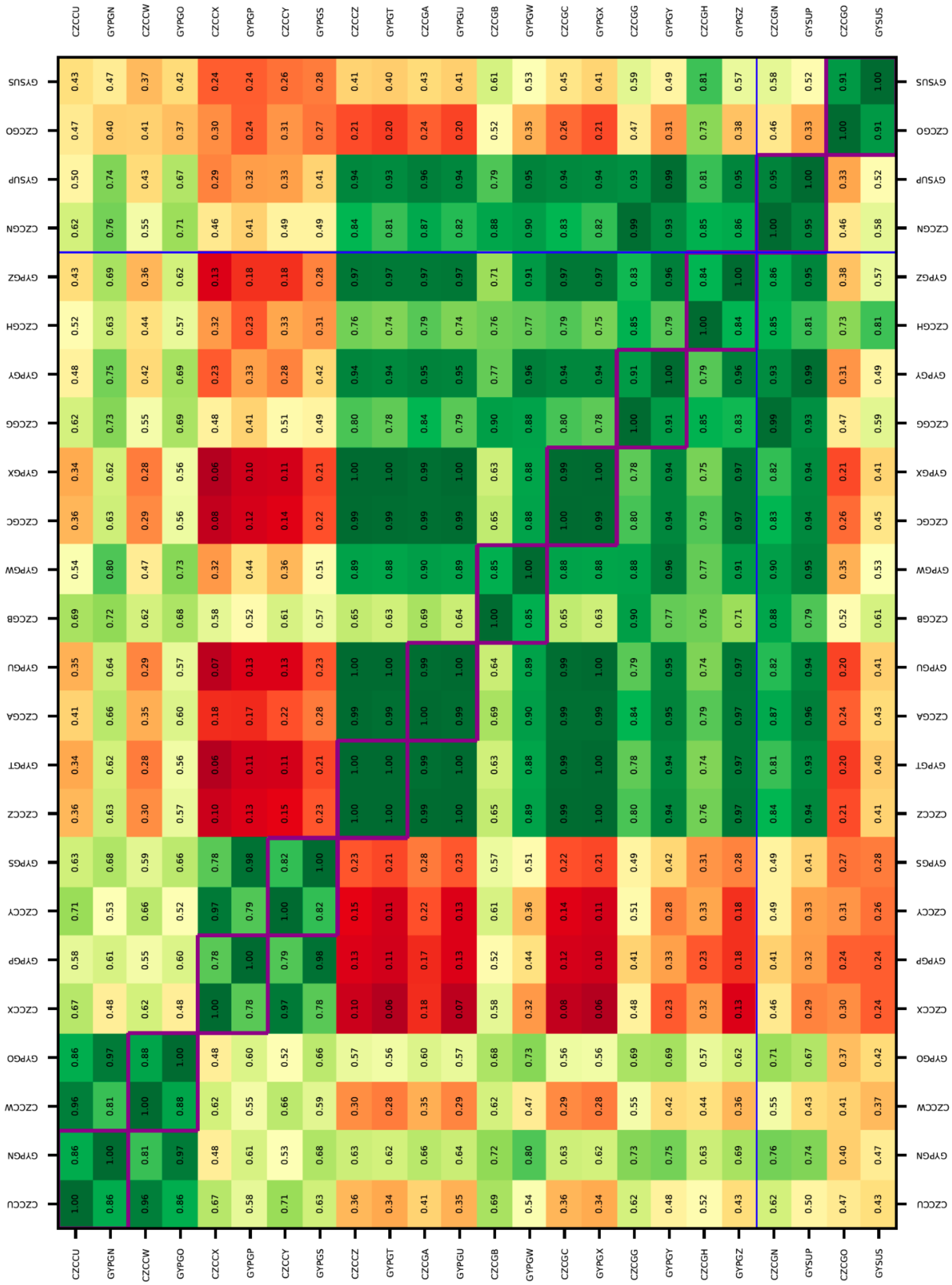

**Figure S5**: Raw gene counts were used to evaluate the level of correlation between biological replicates using Pearson's correlation. Refer to Table S3 for the identity of the sample libraries.

**Table S1**: A comparison of the average nucleotide identity of the twenty eight identified *Corynebacterium glutamicum* isolates with sequence information.

| JGI IMG Genome ID | Strain Name | JGI IMG Genome ID | Strain Name | Average Nucleotide Identity (%) |
|---|---|---|---|---|
| 2821586876 | *C. glutamicum* BRC-JBEI 1.1.2 | 2541047465 | *C. glutamicum* Z188 | 99.9987 |
| 2821586876 | *C. glutamicum* BRC-JBEI 1.1.2 | 2554235421 | *C. glutamicum* SCgG1 | 99.9987 |
| 2821586876 | *C. glutamicum* BRC-JBEI 1.1.2 | 2554235422 | *C. glutamicum* SCgG2 | 99.9987 |
| 2821586876 | *C. glutamicum* BRC-JBEI 1.1.2 | 2534681606 | *C. glutamicum* S9114 | 99.9979 |
| 2821586876 | *C. glutamicum* BRC-JBEI 1.1.2 | 2545824695 | *C. crenatum* MT | 99.9755 |
| 2821586876 | *C. glutamicum* BRC-JBEI 1.1.2 | 2585428016 | *C. crenatum* SYPA 5-5 | 99.9718 |
| 2821586876 | *C. glutamicum* BRC-JBEI 1.1.2 | 640427110 | *C. glutamicum* R | 98.9345 |
| 2821586876 | *C. glutamicum* BRC-JBEI 1.1.2 | 2636415695 | *Brevibacterium flavum* ATCC 15168 | 97.8554 |
| 2821586876 | *C. glutamicum* BRC-JBEI 1.1.2 | 2848153583 | *C. glutamicum* ATCC 14067 | 97.8528 |
| 2821586876 | *C. glutamicum* BRC-JBEI 1.1.2 | 2687453329 | *C. glutamicum* YI | 97.8298 |
| 2821586876 | *C. glutamicum* BRC-JBEI 1.1.2 | 2531839299 | *C. glutamicum* ATCC 14067 | 97.8291 |
| 2821586876 | *C. glutamicum* BRC-JBEI 1.1.2 | 2654587970 | *C. glutamicum* B253 | 97.8238 |
| 2821586876 | *C. glutamicum* BRC-JBEI 1.1.2 | 2869752021 | *C. glutamicum* AJ1511 | 97.7627 |
| 2821586876 | *C. glutamicum* BRC-JBEI 1.1.2 | 2859560989 | *C. glutamicum* WM001 | 97.7553 |
| 2821586876 | *C. glutamicum* | 2718218481 | *C. glutamicum* | 97.7534 |

| | | | | |
|---|---|---|---|---|
| | BRC-JBEI 1.1.2 | | ATCC 13869 | |
| 2821586876 | *C. glutamicum* BRC-JBEI 1.1.2 | 2718217803 | *C. glutamicum* XV | 97.7402 |
| 2821586876 | *C. glutamicum* BRC-JBEI 1.1.2 | 2684623129 | *C. glutamicum* CP | 97.7373 |
| 2821586876 | *C. glutamicum* BRC-JBEI 1.1.2 | 2859553545 | *C. glutamicum* B414 | 97.7355 |
| 2821586876 | *C. glutamicum* BRC-JBEI 1.1.2 | 2802429473 | *C. flavum* ZL-1 | 97.7246 |
| 2821586876 | *C. glutamicum* BRC-JBEI 1.1.2 | 2718218485 | *C. glutamicum* ATCC 21831 | 97.6605 |
| 2821586876 | *C. glutamicum* BRC-JBEI 1.1.2 | 2850571192 | *C. glutamicum* AR1 | 97.6587 |
| 2821586876 | *C. glutamicum* BRC-JBEI 1.1.2 | 2823997243 | *C. glutamicum* ATCC 13287 | 97.6561 |
| 2821586876 | *C. glutamicum* BRC-JBEI 1.1.2 | 2859564202 | *C. glutamicum* ATCC 13032 | 97.6535 |
| 2821586876 | *C. glutamicum* BRC-JBEI 1.1.2 | 639279306 | *C. glutamicum* Kalinowski ATCC 13032 | 97.6478 |
| 2821586876 | *C. glutamicum* BRC-JBEI 1.1.2 | 2554235405 | *C. glutamicum* MB001 | 97.6418 |
| 2821586876 | *C. glutamicum* BRC-JBEI 1.1.2 | 639279307 | *C. glutamicum* Nakagawa ATCC 13032 | 97.6392 |
| 2821586876 | *C. glutamicum* BRC-JBEI 1.1.2 | 2554235360 | *C. glutamicum* K051 | 97.6305 |
| 2821586876 | *C. glutamicum* BRC-JBEI 1.1.2 | 2687453317 | *C. glutamicum* USDA-ARS-USMA RC-56828 | 97.3362 |

**Table S2: Strain and Plasmids Used in this Study.**

| Strain/Plasmid Name | Genotype | Reference |
|---|---|---|
| JBEI-18084 | *C. glutamicum* ATCC 13032 Δ*cglIM* Δ*cgLIR* Δ*cgLIIR* biotin auxotroph, NxR. | A kind gift from Sarah Richardson. Described in Baumgart *AEM*, 2013. [Sequenced at JGI. ProjectID: 503247] |
| JBEI-7936 | *C. glutamicum* BRC-JBEI 1.1.2 biotin auxotroph, NxR | Sasaki and Eng *et al,* BfB, 2019. [Sequenced at JGI. ProjectID: Ga0373873] |
| JBEI-19566 | *C. glutamicum* BRC-JBEI 1.1.2 Δ*ldh* Δ*idsA* biotin auxotroph, NxR derived from JBEI-7936 | Sasaki and Eng *et al,* Met Eng Comm. [Sequenced at JGI. Project ID: 503805] |
| JBEI-19559 | *E. coli* DH10B {pEC-XK99E-AK-IP-bypass} KanR | Sasaki and Eng *et al,* BfB, 2019. |
| JBEI-19632 | *E. coli* DH10B {pEC-XK99E-AK-IP-bypass-*S. pomeroyi* HMGR (*substitution*)} KanR | Sasaki and Eng *et al,* BfB, 2019. |

**Table S3**: RNAseq Sample key and library QC metrics for samples analyzed in the fed-batch campaign.

| **Library Name** | **Sample Id** | **Raw Reads** | **Filtered Reads** | **Sample Name** |
|---|---|---|---|---|
| CZCCU | 191641 | 6565438 | 2547302 | shake flask / final seed train culture CGXII + 4% glucose |
| GYPGN | 191641 | 3783644 | 3447814 | shake flask / final seed train culture CGXII + 4% glucose |
| CZCCW | 191642 | 10201048 | 4842164 | shake flask / final seed train culture CGXII + 4% glucose |
| GYPGO | 191642 | 8921860 | 8194336 | shake flask / final seed train culture CGXII + 4% glucose |
| CZCCX | 191643 | 9942954 | 1265252 | 6.5 hrs post innoc 2 L fed-batch CGXII + 50 mM [Chl][Lys] |
| GYPGP | 191643 | 8646938 | 8023706 | 6.5 hrs post innoc 2 L fed-batch CGXII + 50 mM [Chl][Lys] |
| CZCCY | 191644 | 17316250 | 3273128 | 6.5 hrs post innoc 2 L fed-batch CGXII |
| GYPGS | 191644 | 6786960 | 6327830 | 6.5 hrs post innoc 2 L fed-batch CGXII |
| CZCCZ | 191645 | 10710608 | 7635578 | 16.58 hrs post innoc 2 L fed-batch CGXII + 50 mM [Chl][Lys] |
| GYPGT | 191645 | 7781964 | 5910924 | 16.58 hrs post innoc 2 L fed-batch CGXII + 50 mM [Chl][Lys] |
| CZCGA | 191646 | 5558460 | 499094 | 16.58 hrs post innoc 2 L fed-batch CGXII |
| GYPGU | 191646 | 5876548 | 4501932 | 16.58 hrs post innoc 2 L fed-batch CGXII |
| CZCGB | 191647 | 8645056 | 8167270 | 24 hrs post innoc 2 L fed-batch CGXII 4% + 50 mM [Chl][Lys] |
| GYPGW | 191647 | 8696772 | 7905458 | 24 hrs post innoc 2 L fed-batch CGXII 4% + 50 mM [Chl][Lys] |
| CZCGC | 191648 | 12856598 | 5616520 | 24 hrs post innoc 2 L fed-batch CGXII |
| GYPGX | 191648 | 6564560 | 5652508 | 24 hrs post innoc 2 L fed-batch CGXII |
| CZCGG | 191649 | 8383158 | 7336994 | 41 hrs post innoc 2 L fed-batch CGXII + 50 mM [Chl][Lys] |
| GYPGY | 191649 | 8162496 | 7448506 | 41 hrs post innoc 2 L fed-batch CGXII + 50 mM [Chl][Lys] |
| CZCGH | 191650 | 6516938 | 784488 | 41 hrs post innoc 2 L fed-batch CGXII |
| GYPGZ | 191650 | 7872554 | 6521852 | 41 hrs post innoc 2 L fed-batch CGXII |
| CZCGN | 191651 | 6702200 | 868060 | 65 hrs post innoc 2 L fed-batch CGXII + 50 mM [Chl][Lys] |
| GYSUP | 191651 | 7061322 | 310186 | 65 hrs post innoc 2 L fed-batch CGXII + 50 mM [Chl][Lys] |
| CZCGO | 191652 | 19711092 | 1380996 | 65 hrs post innoc 2 L fed-batch CGXII |
| GYSUS | 191652 | 7430190 | 90368 | 65 hrs post innoc 2 L fed-batch CGXII |
